# Variation of the *Rdr1* gene insertion in wild populations of *Nicotiana benthamiana* (Solanaceae) and insights into recent species divergence

**DOI:** 10.1101/2022.02.01.478068

**Authors:** Luiz A. Cauz-Santos, Steven Dodsworth, Rosabelle Samuel, Maarten J.M. Christenhusz, Denise Patel, Taiwo Shittu, Aljaž Jakob, Ovidiu Paun, Mark W. Chase

## Abstract

- One of the most commonly encountered and frequently cited laboratory organisms worldwide is classified taxonomically as *Nicotiana benthamiana* (Solanaceae), an accession of which, typically referred to as LAB, is renowned for its unique susceptibility to a wide range of plant viruses and hence capacity to be transformed using a variety of methods. However, the origin and age of LAB and the evolution of *N. benthamiana* across its wide distribution in Australia remains relatively underexplored.
- Here, we have used multispecies coalescent methods on genome-wide single nuclear polymorphisms (SNPs) to assess species limits, phylogenetic relationships and divergence times within *N. benthamiana*.
- Our results show that the previous taxonomic concept of this species in fact comprises five geographically, morphologically and genetically distinct species, one of which includes LAB.
- Remarkably, we provide clear evidence that LAB is closely related to accessions collected further north in the Northern Territory; this species split much earlier from their common ancestor than the other four in this clade and is morphologically the most distinctive. Furthermore, this long-isolated species typically grows in sheltered sites in subtropical/tropical monsoon areas of northern Australia, contradicting the previously advanced hypothesis that this species is an extremophile that has traded viral resistance for precocious development.

## Introduction

*Nicotiana benthamiana* is an important model organism widely employed since the early 1950s in research on plant/virus interactions thanks to its unusual susceptibility to a wide range of viruses (Quacquarelli & Avgelis, 1975; Goodin *et al*., 2008) and more recently production of pharmaceuticals (van Zyl *et al*., 2016; Reed & Osbourn, 2018). This species, widely distributed in the tropical and subtropical regions of Australia, is a member of *Nicotiana* section *Suaveolentes*, an allopolyploid group that originated in South America around 5-6 Mya (Clarkson *et al*., 2017a; Schiavinato *et al*., 2020). This clade colonized the arid interior of Australia only in the last two million years, where the plethora of species (c. 50), many of these newly described are commonly encountered (Chase *et al*., 2018, 2021a).

The species of *Nicotiana* sect. *Suaveolentes* occur throughout Australia (but not in Tasmania), and *N. benthamiana* typically grows in sheltered sites on the south sides of rocky outcrops and gorges where it is protected from the extremes of heat and seasonal drought typical of these regions (Chase *et al*., 2018). Despite this preference for sheltered sites, *N. benthamiana* has been labelled an “extremophile” adapted to hot, dry habitats, and it was hypothesized that it has “traded” resistance to viruses for important adaptations, especially precocious flowering and larger seeds (presumably containing more resources), to extreme environmental conditions (Bally *et al*., 2015). Its susceptibility to a large number of plant viruses is a result of loss of function of the RNA dependent RNA polymerase 1 (*Rdr1*) gene caused by an insertion of 72 bp, and a single specific genotype (LAB) harbouring this insertion has been used widely as a model in plant biology (Yang *et al*., 2004; Wylie *et al*., 2015; Bally *et al*., 2015, 2018). LAB has been documented to have originated from a 1936 collection of seeds made at the Granites Goldmine in the Tanami Desert of the Northern Territory and sent to the University of California, from where it made its way into many laboratories after its susceptibility to a wide range of plant viruses was realized (Bally *et al*., 2015). Although genetic evidence pointed to a relationship of LAB with a South Australian accession (putatively from the Everard Ranges in northern South Australia) in a previous study (Bally *et al*., 2015), *N. benthamiana* is in fact undocumented by any vouchered collections from this area (Australian Virtual Herbarium; AVH.chah.org.au/; Fig. 1). Thus, we doubt the Everard Ranges provenance is accurate and conclude that this material of *N. benthamiana* was a contaminant of unknown origin (but from somewhere else in Australia). Apart from the still-open questions about the origin and age of LAB, the evolution of *N. benthamiana* across its wide distribution in Australia remains relatively underexplored.

**Fig. 1:**
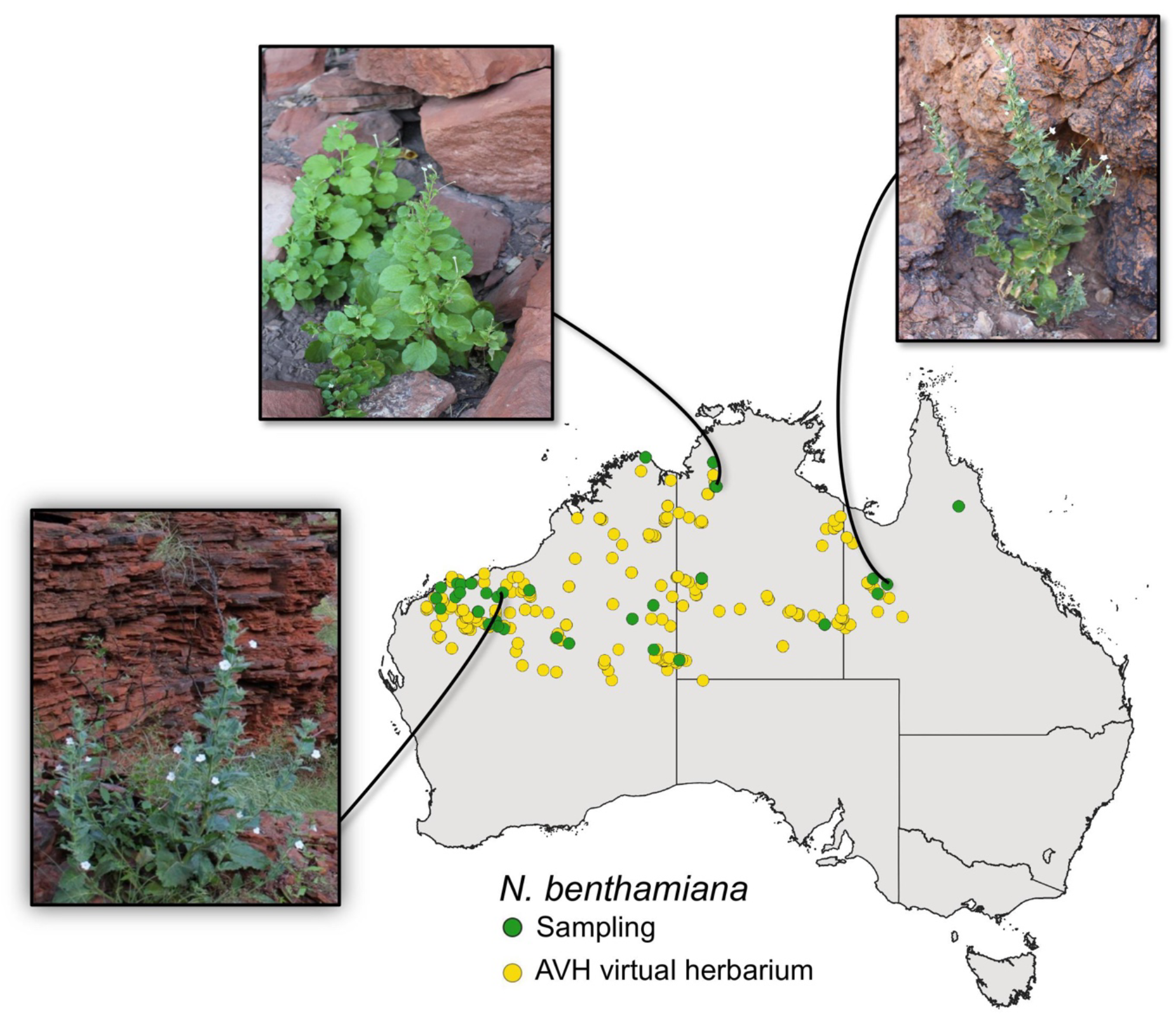
Map showing the distribution of *N. benthamiana* and the sampling locations (green dots) for the accessions in this study. The distribution of *N. benthamiana* in Australia (yellow dots) was based on data from the Australasian Virtual Herbarium (AVH). Note the absence of accessions from the north-central part of South Australia, where the purported SA sample (Bally *et al*. 2015) was supposed to have originated. The images of the *N. benthamiana* plants from Western Australia, Northern Territory and Queensland show the ecological habitat for this species growing in sheltered sites on sides of rocky outcrops and gorges. The coordinates and details of the sampled material is provided in Supplementary Table 1.

Burbidge (1960) and Chase & Christenhusz (2018) hypothesized that there is potentially more than one species within the broader concept of *N. benthamiana* based on patterns of morphological variation (also documented in Bally *et al*., 2015). How widespread species like *N. benthamiana* maintain genetic and phenotypic cohesion across multiple different climatic zones and pronounced ecogeographic barriers in the face of fragmentation and population divergence has been much debated (Rosenzweig, 1995; Pigot *et al*., 2012). Such populations are predisposed to diverge due to a combination of drift and local adaptation (Irwin, 2002; Schluter & Conte, 2009; Barraclough, 2019). Many widely distributed species are also artifacts of the piecemeal delimitation that afflicts the taxonomic process (i.e., different authors treating different sets of species at different times), which is likely to conceal cryptic diversity in such species.

Here, we have used multispecies coalescent methods on genome-wide single nuclear polymorphisms (SNPs) to assess species limits and phylogenetic relationships within *N. benthamiana*, utilizing vouchered specimens from across its natural range (Fig. 1). We document that the former taxonomic concept comprises five genetically distinct species with different morphologies and discrete, allopatric distributions in Australia. We reveal that LAB is closely related to a set of accessions found further north in the Northern Territory and adjacent parts of Western Australia (the Kimberley). Additionally, we find the *Rdr1* insertion to be variable in these Northern Territory populations (including heterozygotes), several of which come from permanently wet (spring fed) sites that also host ferns and *Livistona* palms. Our results strongly contradict the hypothesis of the *Rdr1* mutation as adaptative in arid environments and the characterization of *N. benthamiana* as an extremophile (Bally *et al*., 2015).

## Materials and Methods

### Plant material

For this study, the sampling included 44 individuals, of which 36 are from *N. benthamiana* (11 from viable seeds recovered from herbarium specimens up to 27 years old), four from *N. karijini* and four from *N. gascoynica*, these last two species closely related to *N. benthamiana* and serving as outgroups (Supplementary Table 1). Most samples were collected by Chase and Christenhusz in the wild and are vouchered in three major Australian Herbaria (BRIS, DNA and PERTH, the standard acronyms for these collections). The *N. benthamiana* sampling covers the distribution of this species in Australia, as previously accepted, with accessions from Queensland, Northern Territory and Western Australia. Additionally, the *N. benthamiana* lab strain (TW16; LAB) obtained from the U.S. Department of Agriculture (USDA) was also included. The maps with the sampling locations were generated in QGIS v. 3.20.3 (QGIS Development Team, 2021), the map layer with the vectors from the drainage and rivers divisions were obtained from the Australian Government Bureau of Meteorology (http://www.bom.gov.au/water/geofabric/). The provenance data (latitude, longitude) of the accessions used to construct the map of the *N. benthamiana* distribution were obtained from the Australasian virtual herbarium (https://avh.chah.org.au).

### Collecting and import permits

All field-collected material is covered under the following collecting permits to MWC and MJMC: Western Australia SW017148, CE006044, Northern Territory 58658, and Queensland PTU-18001061. Permission to remove seeds from herbarium specimens was obtained from the curators/collections managers of the following herbaria: BRI, NT, and PERTH. All seeds were imported into the UK following published guidelines, and plants were grown in quarantine at the Royal Botanic Gardens, Kew, UK import permit DEFRA PHL2149/194627/5NIRU CERT:106-2019; HMRC TARIFF CODE: 0601209090.

### DNA extraction, library preparation and sequencing

Total genomic DNA was isolated from ca. 20 mg of silica-dried leaf tissue. The material was pre-treated for 20 min in ice-cold sorbitol buffer (100 mM tris-HCl, 5mM EDTA, 0.35 M sorbitol, pH 8.0), and the extraction was conducted under the cetyltrimethylammonium bromide (CTAB) procedure. Subsequently, DNA was purified using the NucleoSpin gDNA clean-up Kit (Machery-Nagel, Germany), according to the manufacturer’s instructions. DNA content was quantified using a Qubit 3.0 Fluorometer and the dsDNA HS Assay Kit (Thermo Fisher Scientific, Waltham, MA, USA).

The RADseq libraries were prepared by single digestion according to a protocol established in previous studies (Paun *et al*., 2016). The DNA samples were digested with the PstI enzyme, and samples were processed in batches of individuals using inline and index barcodes that were distinct from one another by at least three nucleotide positions. The libraries have been sequenced at the VBCF NGS Unit (www.vbcf.ac.at/ngs) on an Illumina HiSeq 2500 as pair-end reads of 125 bp.

### SNP calling from RADseq data

Raw reads were first processed in BamIndexDecoder v.1.03 (included in Picard Illumina2Bam package, available from http://gq1.github.io/illumina2bam/) and demultiplexed in sublibraries according to index barcodes. Then, a demultiplexing in individuals was conducted in process_radtags from Stacks v.1.47 (Catchen *et al*., 2013) using the inline barcodes, in conjunction with a quality filtering that removed reads containing any uncalled base and those with low quality scores, but rescued barcodes and cut sites with maximum one mismatch.

The reads were mapped to the reference genome of *N. benthamiana* v.1.0.1 (Bombarely *et al*., 2012) with BWA MEM v. 0.7.17 (Li & Durbin, 2009). The option –M was applied during mapping to flag shorter split hits as secondary, and the individual mapping rates were investigated to test for mapping bias, potentially driven by phylogenetic relatedness to the reference individual. The resulted aligned sam file was sorted by reference coordinates, and read groups were added using Picard Toolkit (available from http://broadinstitute.github.io/picard/). To improve the mapping quality, a realignment around indels was performed with the Genome Analysis Toolkit v.3.8 (McKenna *et al*., 2010), thinning the data to a maximum of 100,000 reads per interval.

The variants were called in GATK following the best-practices recommendations for DNAseq. First, the GVCF mode of HaplotypeCaller was used to generate an intermediate gVCF file for each sample, and subsequently, all samples were processed in a joint genotyping analysis under the GenotypeGVCFs module. After using VCFtools v.0.1.15 (Danecek *et al*., 2011) to retain only variants presents in at least 50% of the individuals, the vcf file was filtered in the VariantFiltration module of GATK using the following criteria: (1) depth of coverage (DP) < 500; (2) variant confidence (QUAL) < 30.00; (3) variant confidence divided by the unfiltered depth (QD) < 2; (4) Phred-scaled P-value for the Fisher’s exact test to detect strand bias (FS) > 60; (5) a root mean square of mapping quality across all samples (MQ) < 40; (6) u-based z-approximation from the rank sum test for mapping qualities (ReadPosRankSum) < −8.0; and (7) u-based z-approximation from the rank sum test for the distance from the end of the reads with the alternate allele (MQRankSum) < −12.5.

After SNP calling and filtering first in GATK, we retained a total of 3,283,370 variable sites. These data were then submitted to an additional filtering in VCFtools to retain only SNPs with a minor allele frequency ≥ 0.066 (i.e., present in at least four haplotypes) and an average depth above 20. Finally, to avoid further use of any pooled paralogs, we used the populations pipeline from Stacks to retain only variable positions with a maximum observed heterozygosity of 0.65, and VCFtools for a final filtering by missing data (allowing a maximum of 20%). The final filtered vcf file contained 46,156 SNPs retained for the *Nicotiana* species analysed in this study.

### Phylogenomic analysis

For estimate the phylogenetic relationships among the accessions, the final filtered vcf file was converted to a PHYLIP format using PGDspider v.2.1.1.0 (Lischer & Excoffier, 2012) and subsequently the invariant sites were removed with the script ascbias.py (https://github.com/btmartin721/raxml_ascbias). After removing invariant sites, 14,818 SNPs were retained and used for the phylogenetic reconstruction. The maximum likelihood analyses were performed in RAxML v.8.2.12 (Stamatakis, 2014) with the PHYLIP file containing concatenated SNPs using a recommended correction of the likelihood for ascertainment bias (Lewis, 2001). The analysis was conducted under the general time reversible model of nucleotide substitution and the CAT approximation of rate heterogeneity (GTRCAT), searching for the best-scoring ML tree with 1,000 rapid bootstrap replicates. The best tree was then visualized and annotated in FigTree v.1.4.4 (http://tree.bio.ed.ac.uk/software/figtree/).

### Admixture and genetic clustering analyses

We identified genetic groups and evidence of hybridization/introgression by construction of a coancestry heatmap and admixture proportions for the 36 accessions of the *N. benthamiana* group. For this purpose, we used the indel-realigned .bam files to calculate genotype likelihoods based a genotype-free method implemented in ANGSD v.0.930 (Korneliussen *et al*., 2014). In the final set, we retained only sites with data for at least 50% of individuals and with a minimum 20 base and mapping quality. For 1,253,966 high-confidence (p < 1e-6) variable positions that had a minor allele shared by at least three individuals, we inferred the major and minor alleles frequencies under a GATK-based genotype likelihood model. Starting from covariance matrices calculated using PCangsd (Meisner & Albrechtsen, 2018) from the genotype likelihoods, we further visualized coancestry of the different accessions using the heatmaps.2 function (GPLOTS v.3.0.1.1) (Warnes *et al*., 2020).

We further accessed the population structure among the *N. benthamiana* accessions inferring individual admixture proportions with NGSadmix (Skotte *et al*., 2013) using the genotype likelihoods inferred in ANGSD. First, we used the genotype likelihoods to select unlinked sites with a distance of 10,000 bp between variants, resulting in 46,961 sites. Then, the analysis was conducted using K = 1 to K = 10 as number of clusters for the admixture inference, 20 seeds to initialize the EM algorithm, and only variants with minor allele frequency above 0.05. The results for the different Ks were evaluated based on the Evanno method (EVANNO *et al*., 2005) available in CLUMPAK (http://clumpak.tau.ac.il/bestK.html), and the final plot was produced in R.

To assess the genomic relatedness of *N. benthamiana* and the closely related species *N. gascoynica* and *N. karijini*, we created an additional genotype likelihoods call in ANGSD for a total of 44 accessions. We used the same parameters as described previously, retaining only sites with data for at least 50% of individuals and with a minimum 20-base mapping quality. We obtained 1,748,048 high-confidence (p < 1e-6) variable positions that had a minor allele shared by at least three individuals, which were used to infer the major and minor alleles frequencies under a GATK-based genotype likelihood model. Finally, we calculated the covariance matrices using pcangsd and produced the coancestry heatmap using the heatmaps.2 function (GPLOTS 3.0.1.1)

We also assessed the historical relationships of the groups of the *N. benthamiana* species complex with *N. gascoynica* and *N. karijini* using TREEMIX 1.12 (Pickrell & Pritchard, 2012). This method produces a maximum likelihood graph linking sampled populations to their most common ancestor including migration edges. For this analysis, we produced the input file using VCF-tools v. 0.1.15 (Danecek *et al*., 2011) and Plink v. 0.70 (Weeks, 2010) to convert the vcf file to a frequency file and used the *plink2treemix* script (https://bitbucket.org/nygcresearch/treemix/downloads) to obtain the final file used in TREEMIX. We performed TREEMIX using *N. gascoynica* as outgroup and tested the number of migrations varying from zero to five, assessing the significance of each added migration edge and the residue covariance matrix to assess the fit of the model to the data. The final results were visualized and illustrated in R.

### Inbreeding and private alleles

Individual-based inbreeding coefficients (*F*) and private alleles were estimated for the *N. benthamiana* accessions using the vcf file containing 46,156 SNPs. The first analysis was conducted in VCFtools v.0.1.15(Danecek *et al*., 2011) using the --*het* function, which calculates the measure of heterozygosity per-individual, and plotting of the results was performed in R. The number of private alleles per group was calculated for the SNP data using the populations routine from Stacks v.1.47 (Catchen *et al*., 2013).

### Species delimitation and species tree

To perform Bayesian species delimitation, a smaller dataset was created in VCFtools v.0.1.15 using the filtered vcf file to select unlinked biallelic SNPs (> 10,000 bp distance on the same contig) not allowing for any missing data per locus. For the species delimitation, we constructed a matrix including a total of 25 individuals (22 *N. benthamiana* accessions not exhibiting admixture and three *N. gascoynica*) with at least three individuals per each group identified in the models, which included 10,443 unlinked single nucleotide polymorphisms (SNPs). Taking into account the results from the coancestry heatmaps and TreeMix, we did not include accessions of *N. karijini* in the species delimitation and species tree analysis.

The vcf file containing unlinked SNPs were converted to PHYLIP and then to NEXUS format using PGDSpider v.2.1.1.0, and the input XML files were created using BEAUti v.2.4.8 (Bouckaert *et al*., 2014) and edited to a path-sampling analysis. We tested ten models for the *N. benthamiana* accessions and used *N. gascoynica* as outgroup. The first model was the current taxonomic concept for *N. benthamiana* as single species, and then subsequently splitting in up to six groups (Supplementary Fig. 1). The Bayesian species delimitation analyses were performed in SNAPP v.1.2.5 (Bryant *et al*., 2012) and the marginal likelihood was calculated using 24 initialization steps and 1 million chain-lengths for each model, with data stored every 1,000 generation and excluding 10% of these as burn in. The optimal model of species assignment was defined by calculating the Bayes factor based on the marginal likelihood values.

We used SNAPP v.1.2.5 to construct a coalescent species tree using the optimal model defined in the species delimitation analyses. This analysis was conducted with the same dataset of unlinked SNPs, with 15 million as the chain-length and saving a tree every 1,000th generation. Convergence of the run was monitored with the ESS values from the log-file with Tracer v.1.6 (Rambaut *et al*., 2018). After removing the first 10% of trees as burn in, the SNAPP trees were visualized as a cloudogram using Densitree (Bouckaert & Heled, 2018), and the values of posterior probabilities for each clade were calculated using Treeannotator v.1.8.3 (Drummond *et al*., 2012). The species tree was calibrated using 5e-09 as rate of substitution per site per generation (Schiavinato *et al*., 2020) and one year as generation time for the *N. benthamiana* group and outgroup (these plants are nearly always annuals and rarely live more than a single year). To obtain the divergence times, we rescaled the results using the total length of investigated sites within the loci included and total number of polymorphic sites across their length.

### *Rdr1* amplification and sequencing

Samples for all *N. benthamiana* accessions were screened for the presence of the 72 bp insertion in the *Rdr1* gene. Total DNA was extracted from approximately 10-20 mg of silica-dried leaf tissue, using the Qiagen DNeasy Plant Mini Kit (Qiagen, Germany), according to the manufacturer’s instructions. Initial PCR screening used the rdr1 Fwd and rdr1 Rev primers described previously in Bally *et al*. (2015). These were subsequently redesigned due to variation in amplification efficiency across samples that could not be improved by altering PCR conditions (e.g., annealing temperature). The following primers were used to amplify a smaller product (~570-650 bp) spanning the insertion site, to further screen all populations for the presence of the insert: RDR1SDF (5’ – 3’)— TGCATCAAGACCTGGCCTTA; RDR1SDR (5’ – 3’)—AGCTTCCTGTGCCTTCAAAG.

To confirm presence, absence or heterozygosity of the *Rdr1* insertion, PCR products were run on a 1.5% agarose gel, at 40 V for approximately 150 minutes, and images subsequently inverted. Products for populations showing either the presence of the insert, heterozygosity, or a close phylogenetic relationship to the LAB accession (i.e., in the NT clade) were sent for confirmation by Sanger sequencing. PCR products were purified using a QIAquick PCR Purification Kit (Qiagen, Germany) or NucleoSpin gDNA clean-up Kit (Machery-Nagel, Germany), according to the manufacturer’s instructions, and sent for sequencing at Eurofins Genomics (Ebersberg, Germany). Chromatograms were visually inspected, and forward and reverse reads were assembled de novo in Geneious Prime v.2021.0.3.

### Seed germination, time to flowering and capsule maturity

Seed sizes were measured for mature, dry seeds, stored for a minimum of one year at 5 C. The other traits were determined by growing plants in a temperature-controlled greenhouse, in which night and day temperatures were 18 C and 24 C, respectively. Growing medium, watering etc. were uniform for all plants. Time to first flowering was somewhat ambiguous in some accessions because cleistogamous (unopened) flowers were produced before normal (chasmogamous) flowers, and in these cases perianth withering was used to assess flowering date.

## Results

### Phylogenomic relationships in *N. benthamiana*

In our study, we investigated the evolution of the putative *N. benthamiana* species complex in Australia by producing restriction site-associated DNA sequencing data (RADseq) (Baird *et al*., 2008) that was analysed with respect to the available reference genome of *N. benthamiana* v.1.0.1 (Bombarely *et al*., 2012). A maximum likelihood analysis (RAxML) identified *Nicotiana karijini* as sister to the *N. benthamiana* complex, within which the 36 accessions were distributed in four well-supported clades (BP 100; Supplementary Fig. 2): i) northern-most Kimberly of Western Australia and the Northern Territory (NT) within which LAB is closely related to accessions from Judbarra National Park and the Fish River Station, areas with permanent springs that are subjected to seasonal monsoon conditions; ii) accessions from the Little Sandy, Gibson and Great Sandy Deserts of eastern Western Australia (eWA); iii) accessions from Queensland and eastern Northern Territory (NT); and iv) the coastal region of north-western Western Australian (WA1). Accessions confined to the larger Pilbara Craton (WA2) including the Hamersley Range were scattered across the tree, with relationships between them and the rest unresolved and weakly supported by the maximum likelihood approach.

### Genetic structure and detection of hybridization

The genetic structure within *N. benthamiana* was explored by inferring population structure and individual admixture proportions using NGSadmix, a method based on genotype likelihoods that can detect recent admixture (Skotte *et al*., 2013). The estimation of the likelihood resulted in K = 2 as the best model evaluated in accordance with the Evanno method (EVANNO *et al*., 2005) (Supplementary Fig. 3), with secondary peaks in deltaK for K = 3 and K = 5 (Supplementary Fig. 4). The K = 2 scenario separates the NT group discussed above (including LAB) from the rest of accessions. The K = 5 clustering corroborates the RAxML and multispecies coalescent results (see below) and supports the division of *N. benthamiana* accessions into five geographically distinct groups described above: NT, QLD, eWA, WA1 and WA2 (Fig. 2a). Although the LAB genotype originated at a site in the Tanami Desert, it displays clear relatedness to the other accessions from further north in the Northern Territory rather than further south in the arid regions, suggesting the LAB accession is an outlier in its species in provenance terms.

**Fig. 2:**
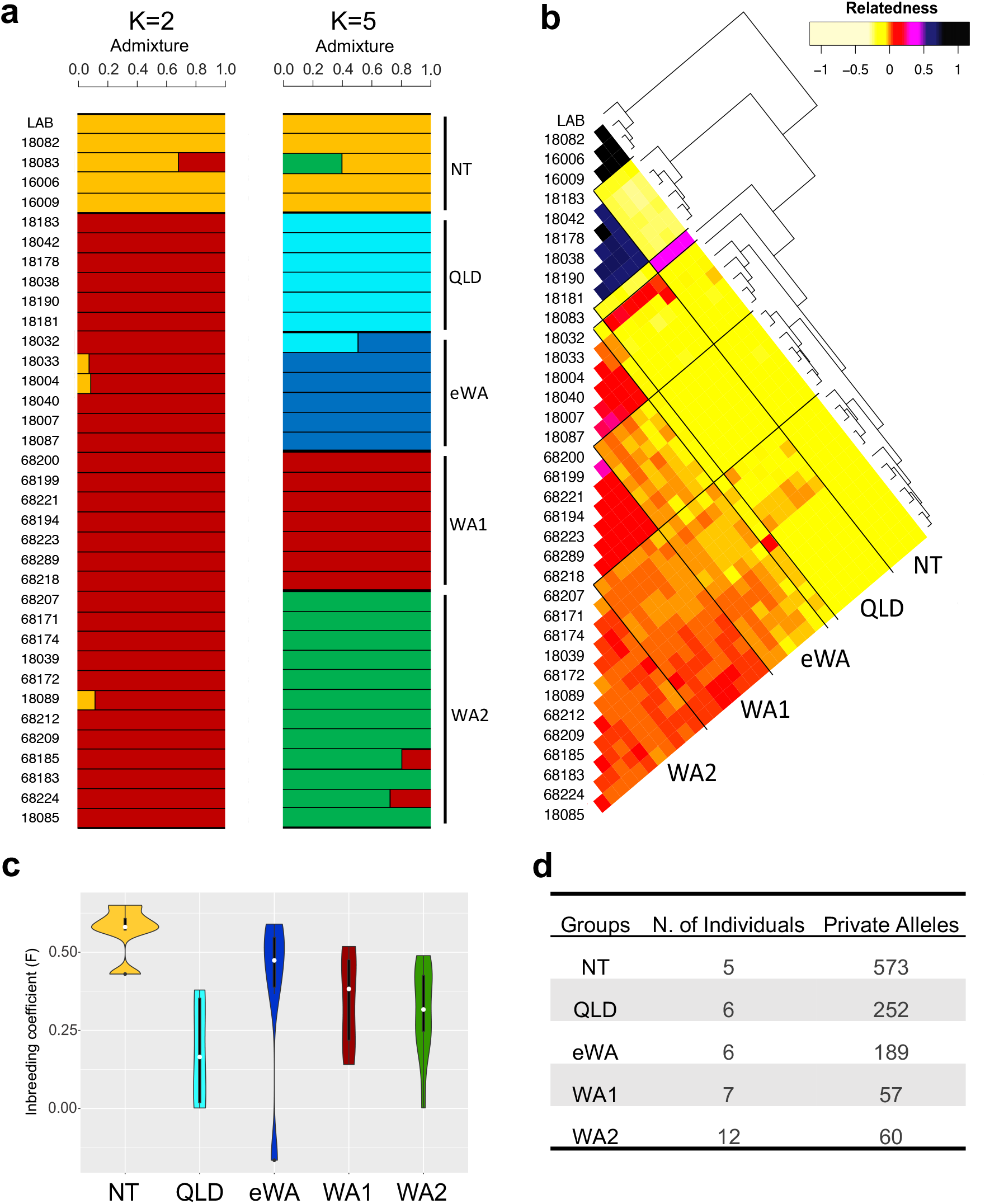
Genetic patterns within *Nicotiana benthamiana*. a) Admixture plot within the *N. benthamiana* accessions for the proposed genetic clusters (K = 2 and K= 5) obtained with NGSadmix. b) Coancestry heatmap of *N. benthamiana* groups constructed based on genotype likelihoods. Darker tones represent higher pairwise relatedness according to legend; estimates for the relationship of one individual to itself have been excluded; note: the order of accessions is different in 2a and 2b because the underlying coancestry tree moved some of the admixed accessions (e.g. 18083) to a different positions. c) Individual-based inbreeding coefficient (F) estimated for the five groups in the *N. benthamiana* complex. d) Private alleles obtained for the *N. benthamiana* groups.

The *N. benthamiana* coancestry heatmap resulted in the identification of introgression in some NT, QLD and eWA accessions (Fig. 2b). These results complement the groupings observed from the genetic clusters obtained in NGSadmix and exhibit an unambiguous structure between the groups, with only a few accessions exhibiting admixture. The coancestry heatmap also corroborates the groupings observed in the RAxML tree, and the NT and QLD clusters exhibit high levels of coancestry within their respective groups. Although well supported in the RAxML tree, the eWA and WA1 clusters exhibit higher levels of diversity; the poorly resolved WA2 cluster from the RAxML tree has the lowest level of coancestry between its accessions, likely the result of recent, but perhaps not on-going (Fig. 2a), gene flow with the accessions of eWA and WA1, which are close geographically.

When estimating individual inbreeding coefficients (F) in the *N. benthamiana* group accessions, we observe that the clear structure identified in the previous analyses is supported by the high level of inbreeding in these accessions (Fig. 2c). The F values were particularly high for all NT accessions (F = 0.43–0.65), consistent with the genetic isolation and distinctiveness observed for NT in the coancestry heatmap. The number of private alleles for the distinct geographic clusters of the *N. benthamiana* species complex (Fig. 2d), ranging from 57 to 573, was highest in the NT cluster including LAB. WA1 and WA2 had the lowest (and a similar) number of private alleles, corroborating the lack of clear differentiation observed in the coancestry heatmap.

In exploring the genomic relatedness among accessions of the *N. benthamiana* species complex and the closely related species *N. gascoynica* and *N. karijini*, we identified a clear structure of the three groups comprising each species, including a high level of coancestry in *N. gascoynica* and *N. karijini* (Supplementary Fig. 5). Introgression with *N. karijini* was identified in some accessions from Queensland and eastern Western Australia. One putative F_1_ hybrid and two accessions with high levels of introgression were also identified, the first between *N. benthamiana* (OLD 18178) from Queensland and *N. karijini*, the second from eastern Western Australia (eWA 18032) with the Queensland (QLD) group, and a third in the NT group (18083) from the Kimberley and *N. benthamiana* from the Pilbara region of Western Australia (WA2). We further investigated the signals of introgression identified in *N. benthamiana* inferring events of migration based on a maximum-likelihood approach in TreeMix. Our TreeMix results shows a population graph with a similar topology to that obtained with RAxML but indicating a migration edge between *N. karijini* and the QLD group (Supplementary Fig. 6), revealing putative old gene flow after the split of these two species from their common ancestor. With this evidence of introgression between *N. karijini* and the *N. benthamiana* species group, we removed the *N karijini* accessions from the species delimitation and species tree analyses to avoid problems in these coalescent-based inferences.

### Species limits and divergence times in the *N. benthamiana* complex

Considering the clear structure of the different groups in *N. benthamiana*, we evaluated the hypothesis of potentially more than one biological species within the broader concept of *N. benthamiana* by conducting a coalescent-based, Bayesian species delimitation analysis. The highest marginal likelihood estimates and the Bayes factor test indicated that the best species delimitation model was the one splitting *N. benthamiana* into five species, corresponding to NT, QLD, eWA, and the two groups from the Pilbara region of Western Australia, WA1 and WA2 (Fig. 3a), the latter restricted to the large Pilbara Craton, a pre-tectonic crust (Djokic *et al*., 2017) that is 3.6–2.7 billion years old.

**Fig. 3:**
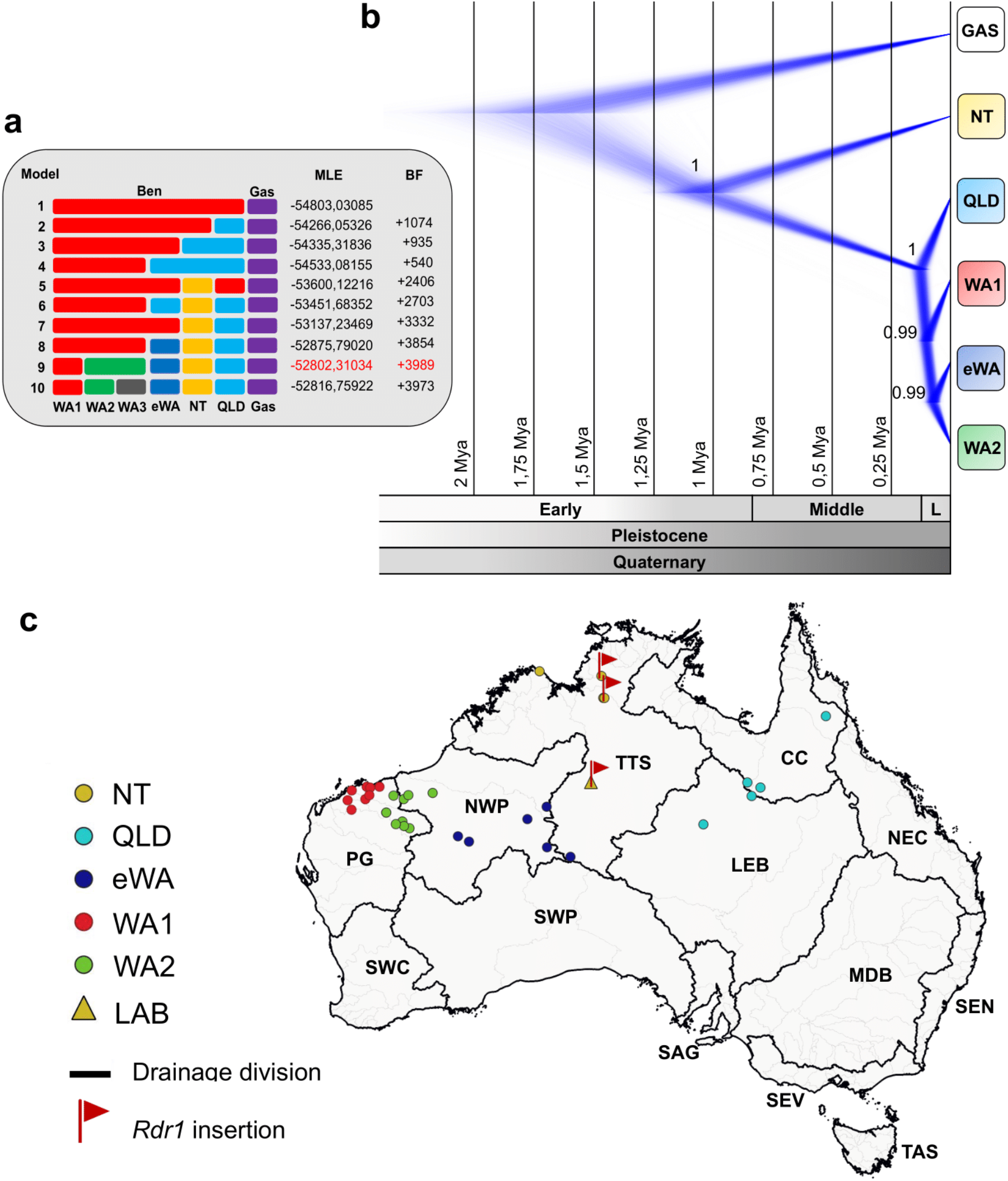
SNAPP multispecies coalescent-based phylogenetic analysis for *Nicotiana benthamiana*. a) Species delimitation models tested for the *N. benthamiana* complex. The best model with the marginal likelihood estimate is highlighted in red. b) Cloudogram from *N. benthamiana* complex species trees with divergence times. L = Late. c) Map of the drainage areas of Australia indicating the sampling locations for the five groups in *N. benthamiana* and highlighting the accessions that contain the 72 bp insertion in the *Rdr1* gene. CC = Carpentaria Coast, LEB = Lake Eyre Basin, MDB = Murray-Darling Basin, NEC = North East Coast (Queensland), NWP = North Western Plateau, PG = Pilbara-Gascoyne, SAG = South Australian Gulf, SEN = South East Coast (NSW), SEV = South East Coast (Victoria), SWC = South West Coast, SWP = South Western Plateau, TAS = Tasmania, TTS = Tanami-Timor Sea Coast.

The SNAPP species tree analysis for the *N. benthamiana* complex resulted in one topology with strong support representing 99.7% of the posterior density distribution. The convergence of the analysis was monitored, and all ESS values were higher than 200. In accordance with the number of private alleles (Fig. 2d), the species tree places NT including LAB as a distant, early splitting relative of the rest, followed by QLD with much more shallow divergences preceding the emergence of WA1, and finally eWA and WA2 as the two most recently diverged groups (Fig. 3b). Divergence time estimation indicated that the diversification of the *N. benthamiana* complex started in the Early Pleistocene, around 1.1 Mya with the split of NT from the rest of the group. However, divergence times for the other four species were more recent and occurred between the end of the Middle and beginning of the Late Pleistocene, around 150,000 years ago (Fig. 3b). When plotting provenance for our sampled accessions on a map of the drainage divisions/rivers of Australia, we found that these five species are largely restricted to one or two adjacent drainage basins (Fig. 3c). The NT group including LAB and the other accessions with the *Rdr1* insertion is restricted to the Tanami-Timor Sea Coast basin.

### Variation in the presence of the *Rdr1* gene insertion

All our results strongly support a close relationship of LAB with these more northerly NT accessions. With respect to the important role of the 72-bp *Rdr1* insertion resulting in virus susceptibility of *N. benthamiana* (and its use as model species), we sought to gain a better understanding of its ecological circumstances and geographical distribution, and we screened for the *Rdr1* insertion in our accessions. PCR amplification and sequencing of all accessions identified two other accessions in the NT group with the *Rdr1* insertion, Victoria River Crossing (Judbarra National Park; *Chase & Christenhusz 16009*) and the Fish River Station (*Cowie 13343, Chase & Christenhusz 18082*) (Fig. 4a). Another accession from Judbarra (at Joe Creek, *Chase & Christenhusz 16006*), 6.7 km from the Victoria Crossing site, did not have the insertion (Figs. 4b and 4c). We had collected leaves from a total of 16 individual plants in these two Judbarra sites (*16006/16007* from Joe Creek and *16008/16009* from Victoria Crossing), from which we amplified the insertion-containing fragment of *Rdr1*, demonstrating that none of the Joe Creek plants has the insert, whereas only one of the Victoria Crossing plants lacked the insert and at least three plants were heterozygous, so the insertion is variable even within some populations (Fig 4b). We had only one plant sampled from the Fish River Station (there was a single plant on the *Cowie 13343* herbarium specimen), which was also heterozygous, so the population from this location must also vary for insertion presence.

**Fig. 4:**
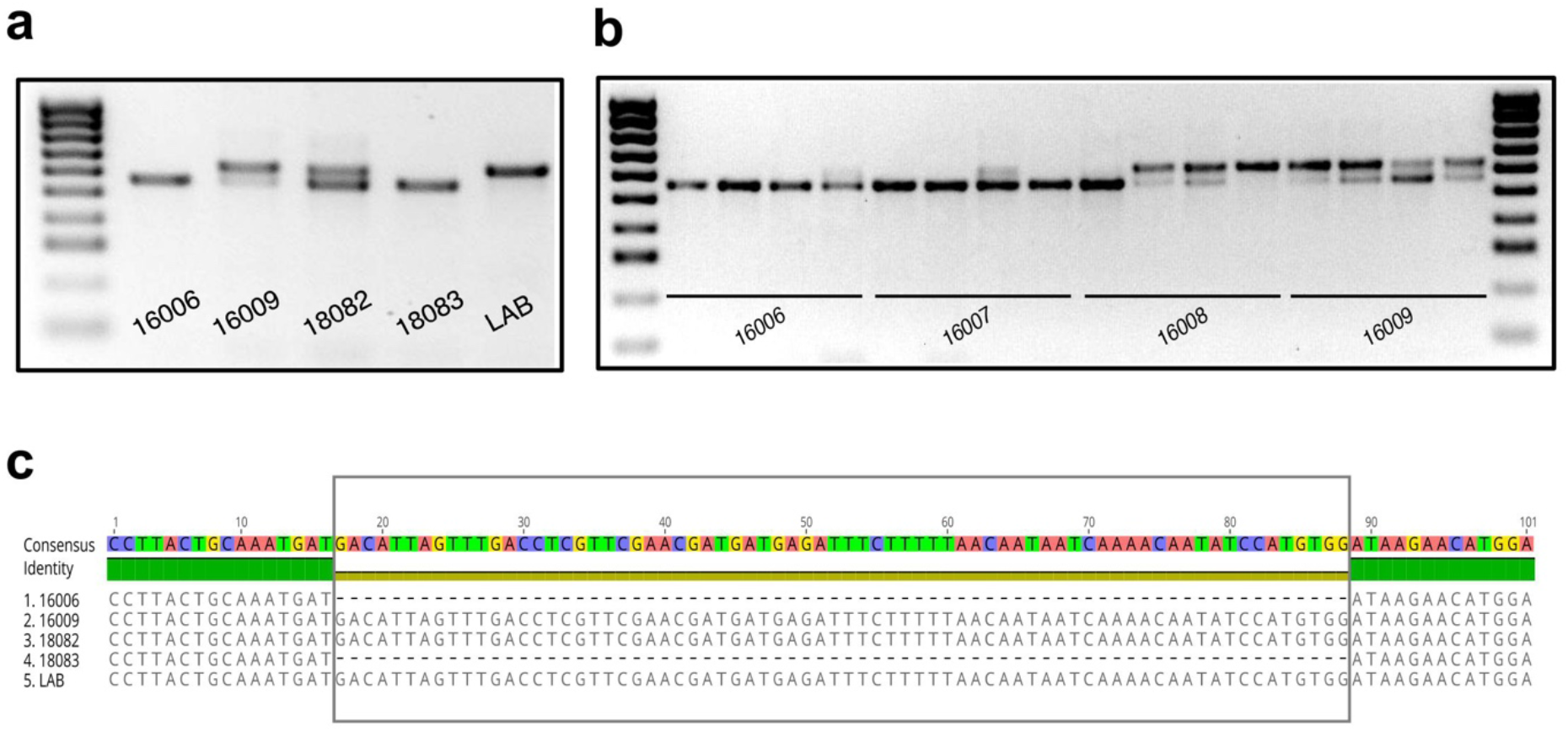
*Rdr1* gene insertion in NT *N. benthamiana* accessions. a) Amplification of the *Rdr1* gene for the individual accessions from Northern Territory included in the RADseq analysis. b) Amplification of the *Rdr1* gene in multiple accessions of populations from northern Northern Territory showing variance in the presence of the 72 bp insertion. c) Alignment of the insertion region from the *Rdr1* gene for the Northern Territory accessions included in the RADseq analysis.

### Germination and flowering times for the *N. benthamiana* accessions

We have kept a log of germination time (summarized in Supplementary Table 2), first flowering, and onset of seed production for all 36 accessions of the *N. benthamiana* group studied here and cannot reproduce the previous results (Bally *et al*., 2015) for the precocious development of LAB. LAB took three days to germinate, but so did insert-lacking *N. simulans* (from highly exposed sites in the extremely arid gibber plains of South Australia) and *N. exigua* (from open to partially shaded sites in eastern New South Wales). The other accessions of the *N. benthamiana* species group followed within the next 1-2 days, and one of the QLD accessions was the first to flower and produce mature fruit. Many other accessions from all areas produced seeds nearly as rapidly as did LAB. Seed size of LAB is larger than many accessions (Bally *et al*., 2015), a trait that is shared with all accessions in which the insertion in *Rdr1* is present (Supplementary Table 2). This same pattern of larger seeds is observed for the two populations from Judbarra National Park, which exhibited heterogeneity for the *Rdr1* insertion. Seed sizes were smaller in the populations lacking the insertion (*16006/16007* from Joe Creek) than in the populations with the insertion (*16008/16009* from Victoria Crossing), and all NT accessions (including LAB) are larger than the other four species in the *N. benthamiana* complex. What seed size contributes to these accessions is unlikely to be related to the “extremophile hypothesis” since it does not appear to contribute to precocious development or shorter time to seed production (Supplementary Table 2).

## Discussion

Our results clearly demonstrate that the previous assumptions (Bally *et al*., 2015, 2018) about the taxonomic status and provenance of LAB were unfounded and have led to a misunderstanding of the ecology of LAB and the context of its viral susceptibility. LAB is clearly not a member of a species widespread across the Australian continent, but rather an isolated, genetically highly derived species confined to the Tanami-Timor Sea coast drainage (principally the Daly and Victoria Rivers in the Northern Territory and the Fitzroy and Ord–Pentecost Rivers in Western Australia). LAB is also not an extremophile species (the Joe Creek and Fish River collections came from places where there are permanent springs, both with abundant ferns and the former with the palm, *Livistona victoriae*), and all sites from where it is known are on the sheltered sides of rock outcrops or cliff faces, so even though some of the plants, e.g. LAB in the Tanami Desert, grow in what is otherwise an arid area it does not have to deal with the extremes of heat and drought these regions experience. The fact that these populations exhibit variation in the presence of the *Rdr1* insert and some are heterozygous for the insertion also raises further questions. Presence of the insertion does affect seed size (Supplementary Table 2), but it does not reduce the time to first flowering and seed release as postulated previously (Bally *et al*., 2015). When the precocious development of LAB demonstrated previously (Bally *et al*., 2015) in a small set of accessions (five plus LAB) is examined in the context of more accessions (here 36), LAB fails to be an outlier. The NT populations (including LAB) exhibit high levels of inbreeding, leading us to conclude that exposure of the populations with the inactivating viral insert to other plants of the NT species is extremely limited and thus few to no opportunities exist in the wild for exchange of viruses, which means that resistance to viruses is unlikely to be detrimental, at least in the short term. Our results thus far reject the notion of it being an extremophile that has traded virus resistance for precocious development, but this is a subject that requires additional research, especially given that the presence of the insertion is correlated with larger seeds.

Another of our novel results is the age of these events. Bally *et al*. (2015) stated that formation of *Nicotiana* sect. *Suaveolentes* took place 20 Mya, based on previous subjective assumptions of the age of this group (Marks *et al*., 2011; Ladiges *et al*., 2011), but molecular clock studies of *N*. sect. *Suaveolentes* have consistently demonstrated that the group is much younger, roughly 6-7 Mya (Clarkson *et al*., 2017a; Schiavinato *et al*., 2020), with the core group of species inhabiting the arid zone of Australia starting to radiate there about 2 Mya. Our results here, developed completely without resort to the previous molecular studies and the methods employed there, corroborate these molecular clock studies (Clarkson *et al*., 2017a; Schiavinato *et al*., 2020). The split between *N. gascoynica* and the *N. benthamiana* species-group at 1.5–2.0 Mya roughly represents the divergence of the core group of species, congruent with the previous estimate of this group (Clarkson *et al*., 2017a). LAB and related accessions diverged much earlier (1.1 Mya) than the other species in the *N. benthamiana* complex, and the timing of the insertion in *Rdr1* would likely be much more recent than this, although we would need a more complete sampling of the populations in this group to obtain a more accurate dating of this event. The remaining species began to diverge only 125,000 ya, with the most recent split between eWA and WA2 occurring only c. 75,000 ya, just before the human occupation of Australia began c. 65,000 ya (Clarkson *et al*., 2017b).

The phylogenetic tree presented in the Bally *et al*. (2015) for the five accessions of *N. benthamiana* (plus LAB) does not agree with our analyses, the result perhaps of the different level of sampling (six versus our 36 accessions). However, their tree is also likely unreliable due to their mixing of diploid and tetraploid accessions/species without separation of paralogs in the two, low-copy nuclear genes used in its production, glutamine synthetase (*Gsl*) and alcohol dehydrogenase C (*AdhC*). For comparison, see Clarkson *et al*. (2015), who also used *Gsl* but separated paralogs such that one set of copies in the allotratrapoloid *N*. sect. *Suaveolentes* fell with the diploid *N. sylvestris* and another copy with other diploids. Failure to separate paralogs if both diploids and allotetraploids are included in their analyses would have unpredictable consequences and make both the tree and the resulting estimates of ages using a molecular clock in Bally *et al*. (2015) totally unreliable.

Finally, *N. benthamiana* as previously circumscribed was considered to be a widespread species with the type of the name (a collection made by Bynoe in 1848, originating from the coast of northwestern Western Australia at Nickol Bay, Karratha (Goodin *et al*., 2008)), and now we have provided evidence that it comprises five species. According to the rule of name priority, the name *N. benthamiana* should be retained for the WA1 group, to which the Bynoe collection, the current holotype, clearly belongs. If nothing were done, this would mean that LAB and its relatives must be recognized as a new species (i.e., with a different name). To avoid the mismatch between this new name and the extensive literature on LAB as *N. benthamiana*, we have proposed conservation of the name with a new type, the original Cleland 1936 collection from the Granites Goldmine, which is preserved in the State Herbarium of South Australia in Adelaide (AD, Australia) (Chase *et al*., 2021b). Thus, we will be proposing new names for the other four species identified in this study, preserving the use of *N. benthamiana* for LAB and preventing confusion in the literature.

## Acknowledgements

We thank Juliane Baar and Daniela Paun for constructing RADseq libraries. Financial support to prepare the RAD library and analyse data were funded by research grants P 26548-B22, P33028-B of the Austrian Science Fund (FWF) and an award (Dr. Anton Oelzelt-Newin Foundation) from the Austrian Academy of Sciences (ÖAW) to Rosabelle Samuel.

## Author contributions

The study was conceived by MWC, RS and OP. MWC and MJMC conducted fieldwork and collected/grew all accessions. LACS, AJ and OP analysed the data. DP and TS performed the amplification, sequencing and analysis of the *Rdr1* insertion under the supervision of SD. MWC and MJMC conducted the inferences of seed germination, time to flowering and capsule maturity. LACS and MWC wrote the manuscript with input from SD, RS and OP.

