## Supplementary material for "Variation of the *Rdr1* gene insertion in wild populations of *Nicotiana benthamiana* (Solanaceae) and insights into recent species divergence"

Supplementary Table 1. Voucher numbers, herbaria and provenance of the plant material.

| Species name | Voucher number (*Chase & Christenhus*)* | Latitude/longitude (S and E; degrees, minutes, seconds) | Provenance (brief locality name, all Australia) |
|---|---|---|---|
| *benthamiana* | *16006* | -15, 36, 22; 131, 5, 3 | Judbarra Gregory NP, Northern Territory |
| *benthamiana* | *16009* | -15, 36, 45; 131, 8, 57 | Judbarra Gregory NP, Northern Territory |
| *benthamiana* | 18004 *Butcher & Albrecht 2048* (PERTH 8855234) | -22, 44, 39; 126, 36, 27 | 115.3 to 118.5 km due West of Kiwirrkurra, Gary Junction Road, Western Australia |
| *benthamiana* | 18007 *Muir 1023* (PERTH 8610134) | -24, 4, 9.75; 123, 9, 58.47 | 400 k E of Capricorn, Moongooloo Rock Hole, Constance Headland, Western Australia |
| *benthamiana* | 18032 *Latz 30766* (NT D0273011); | -24, 24, 18; 127, 45, 58 | Circus Rockhole, Rawlinson Range, Western Australia |
| *benthamiana* | 18033 *Goods 1145* (PERTH 8894183), | -22, 0, 57.3; 127, 44, 38.8 | 4 km S of Bibarrd, Western Australia |
| *benthamiana* | 18038 *Albrecht 12441* (NT D0182096) | -23, 3, 20; 136, 58, 58 | Mt. Tietkens, North Simpson Desert, Northern Territory |
| *benthamiana* | 18039 *Bean 25412* (BRI AQ735848; PERTH 8116067 | -22, 52, 55; 119, 14, 9 | E of Weeli Wolli Creek, 75 km NW of Newman, Western Australia |
| *benthamiana* | 18040 *Latz 22902* (NT D0182743) | -24, 58, 0; 129, 8, 52 | 12 km SE of Docker River Settlement, Learmonth Park, Northern Territory |
| *benthamiana* | 18042 *Wannan 5860* (BRI AQ0855406) | -16, 40, 43; 144, 12, 29 | Beside tributary of Elizabeth Ck, Bellevue, Queensland |
| *benthamiana* | 18082 *Cowie 13343* (CANB 595117.1) | -14, 18, 41; 130, 58, 17 | Fish River Station, Northern Territory |
| *benthamiana* | 18083 *Willing s.n.* (PERTH 8557993) | -14, 2, 12; 127, 19, 31 | King George River tidal estuary, N Kimberley, Western Australia |
| *benthamiana* | 18085 | -22, 11, 35; 116, 14, 14 | Site API-5011. Cardo East, West Pilbara Iron |

| | | | |
|---|---|---|---|
| | *McMaster 25736* (PERTH 8608652) | | project area, Western Australia |
| *benthamiana* | 18087 *Davis 11189* (CANB 698209.1) | -23, 45, 16; 122, 30, 51 | Durba Springs, Canning Stock Route, Western Australia |
| *benthamiana* | *18178* | -20, 53, 7; 140, 20, 59 | Duchess/Cloncurry Road, SSW of Cloncurry, Queensland |
| *benthamiana* | *18181* | -21, 23, 8; 139, 49, 53 | Duchess/Dajarra Road, 5 km southwest of Duchess, Queensland |
| *benthamiana* | *18183* | -21, 6, 47; 139, 48, 54 | Duchess/Mount Isa Road, 55 km southeast of Mount Isa, Queensland |
| *benthamiana* | *18190* | -20, 35, 12; 139, 34, 41 | Lake Moondara Park, Queensland |
| *benthamiana* | *68171* | -23, 17, 3; 119, 39, 30 | Silent Gorge, ca 10 km west of Newman, Western Australia |
| *benthamiana* | *68172* | -23, 9, 5; 119, 20, 5 | Great Northern Highway (95) 11 km from Newman, Western Australia |
| *benthamiana* | *68174* | -23, 2, 27; 118, 51, 4 | Mt Robinson, Western Australia |
| *benthamiana* | *68183* | 22, 21, 30; 118, 17, 4 | Karijini National Park, Hancock Gorge, Western Australia |
| *benthamiana* | *68185* | -22, 21, 25; 118, 17, 13 | Karijini National Park, Weano Gorge, Western Australia |
| *benthamiana* | *68194* | -21, 34, 11; 117, 3, 8 | Millstream-Chichester National Park, Western Australia |
| *benthamiana* | *68199* | -20, 50, 22; 117, 8, 13 | Road from Roebourne to Harding Dam, Western Australia |
| *benthamiana* | *68200* | -20, 53, 19; 117, 20, 26 | ca 2 km WNW of Wittenoom turn off, NW Coastal Highway, Western Australia |
| *benthamiana* | *68207* | -21, 21, 42; 118, 42, 55 | Great Northern Highway, Western Australia |
| *benthamiana* | *68209* | -21, 33, 44; 119, 19, 14 | Woodstock-Marble Bar road, Western Australia |

| | | | |
|---|---|---|---|
| *benthamiana* | *68212* | -21, 19, 54; 119, 35, 28 | Woodstock-Marble Bar road, Western Australia |
| *benthamiana* | *68218* | -20, 50, 16; 117, 53, 21 | NW Coastal Hwy, ca. 5 K E of Whim Creek, Western Australia |
| *benthamiana* | *68221* | 21, 3, 12; 116, 15, 12 | Fortescue River Mouth Road, Western Australia |
| *benthamiana* | *68223* | -21, 38, 4; 116, 0, 35 | Pannawonica Road, Western Australia |
| *benthamiana* | *68224* | -21, 39, 39; 116, 16, 30 | Pannawonica Road, west of Pannawonica, Western Australia |
| *benthamiana* | *68289* | -21, 20, 2; 117, 17, 18 | Millstream-Chichester National Park, Western Australia |
| *benthamiana* | 18089 Chinnock 9599 (AD 170035; NT A0109792; PERTH 7094582) | -21, 12, 13; 121, 1, 21 | Oakover River on Woodie Woodie Rd, Western Australia |
| *benthamiana* LAB | TW16 | -20, 34, 14; 130, 21, 13† | United States Department of Agriculture seed bank |
| *gascoynica* | *68253* | -24, 49, 41; 113, 46, 12 | NW Coastal Hwy, Gascoigne River Crossing, Western Australia |
| *gascoynica* | *68257* | -24, 45, 21; 114, 8, 10 | Gascoyne River, Rocky Pool, Western Australia |
| *gascoynica* | *68265* | -25, 17, 46; 115, 36, 38 | Carnarvon-Mullewa Road, Daurie River crossing, Western Australia |
| *gascoynica* | *68268* | -25, 45, 29; 114, 16, 41 | NW Coastal Hwy, Wooramel River Bridge, Western Australia |
| *karijini* | 18002 | 23, 15, 56; 117, 44, 18.7 | Eastern Range/Channar, Greater Paraburdoo, Pilbara, Western Australia |
| *karijini* | 18009 | 22, 42, 4.7; 117, 23, 58.59 | 1.6 km NE of Mount Turner, Western Australia |
| *karijini* | 18029 | 23, 12, 0; 117, 30, 0 | 21.7 km W of Paraburdoo, Western Australia |

| | | | |
|---|---|---|---|
| *karijini* | 68178 | 22, 23, 25; 118, 16, 3 | Karijini National Park, Joffre Gorge, Western Australia |

*unless otherwise indicated; vouchers deposited at PERTH, DNA or BRIS, depending on the state in which they were collected. †Coordinates for the Granites Goldmine, Northern Territory. Accessions raised from seeds retrieved from herbarium material are documented by secondary vouchers at RBG Kew (K); for these, the original collector, number and herbarium accession number in Australia are also provided.

Supplementary Table 2. *Rdr1* presence/absence, seed sizes and germination, flowering and capsule maturation times for a selection of relevant accessions and species of *N.* sect. *Suaveolentes*. These data were collected for all 36 accessions of the *N. benthamiana* species group studied here, but we do not include all these due to their highly redundant nature. Note: *Nicotiana simulans* and *N. exigua* are distantly related to the *N. benthamiana* complex and do not carry the *Rdr1* insert, but they develop and mature much more quickly compared to the species of the *N. benthamiana* complex.

| **Accession** | **Group** | ***Rdr1* insertion** | **Seed size (microns)** | **Days to germination** | **Days to flowering** | **Days to capsule maturation** |
|---|---|---|---|---|---|---|
| LAB | NT | present | 790–805 | 3 | 55 | 87 |
| *Chase & Christenhusz 16006* | NT | absent | 650–680 | 4 | 53 | 87 |
| *Chase & Christenhusz 16009* | NT | present | 715–755 | 4 | 54 | 94 |
| *Cowie 13343* (18082)* | NT | present | 720-780 | 5 | 54 | 95 |
| *Chase & Christenhusz 18190** | QLD | absent | 530–590 | 4 | 62 | 95 |
| *Chase & Christenhusz 18183** | QLD | absent | 530–585 | 4 | 58 | 95 |
| *Latz 22902 (18040)* | eWA | absent | 510–565 | 5 | 50 | 85 |
| *Goods 1145* (18033)* | eWA | absent | 505–555 | 5 | 64 | 95 |
| *Bean 25412 (18039)* | WA2 | absent | 520–585 | 5 | 60 | 100 |
| *Chase & Christenhusz 68174* | WA2 | absent | 520–580 | 5 | 60 | 102 |
| *Chase & Christenhusz 68199* | WA1 | absent | 530–580 | 5 | 58 | 100 |
| *McMaster 25736 (18085)* | WA1 | absent | 535–585 | 6 | 58 | 102 |
| *N. simulans** | n/a | absent | 580–620 | 3 | 35 | 72 |

| | | | | | | |
|---|---|---|---|---|---|---|
| *N. exigua** | n/a | absent | 450–485 | 3 | 37 | 75 |

*cleistogamous flowers produced before normal (chasmogamous) flowers.

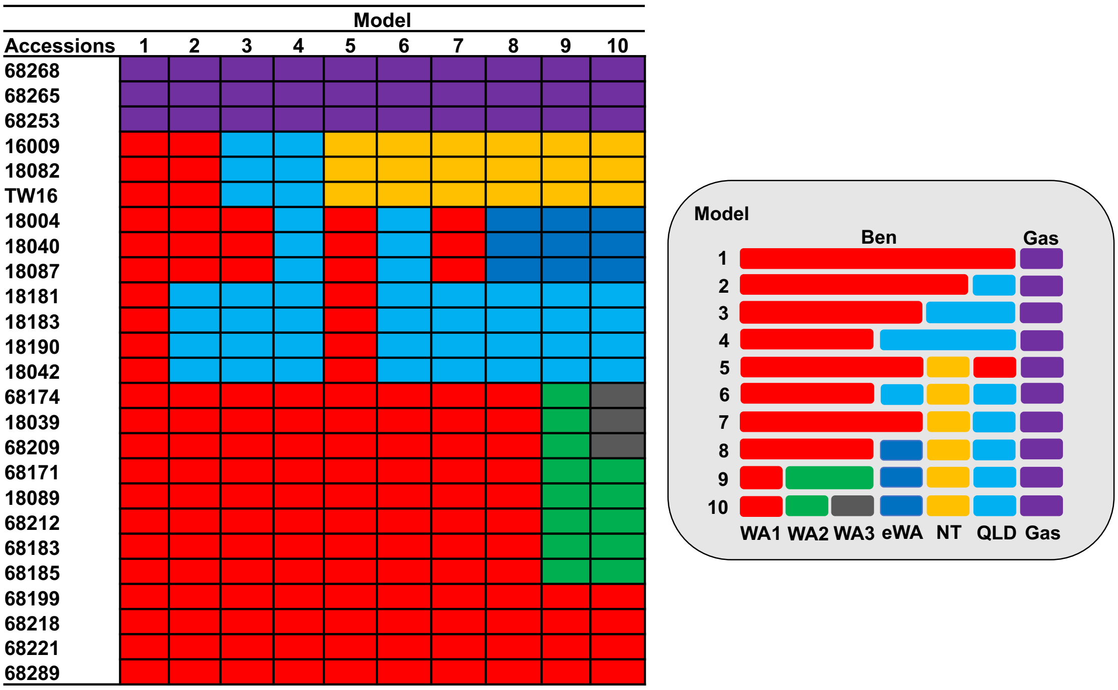


Supplementary Fig. 1. Models and accessions used in the Bayesian species delimitation analysis in SNAPP.

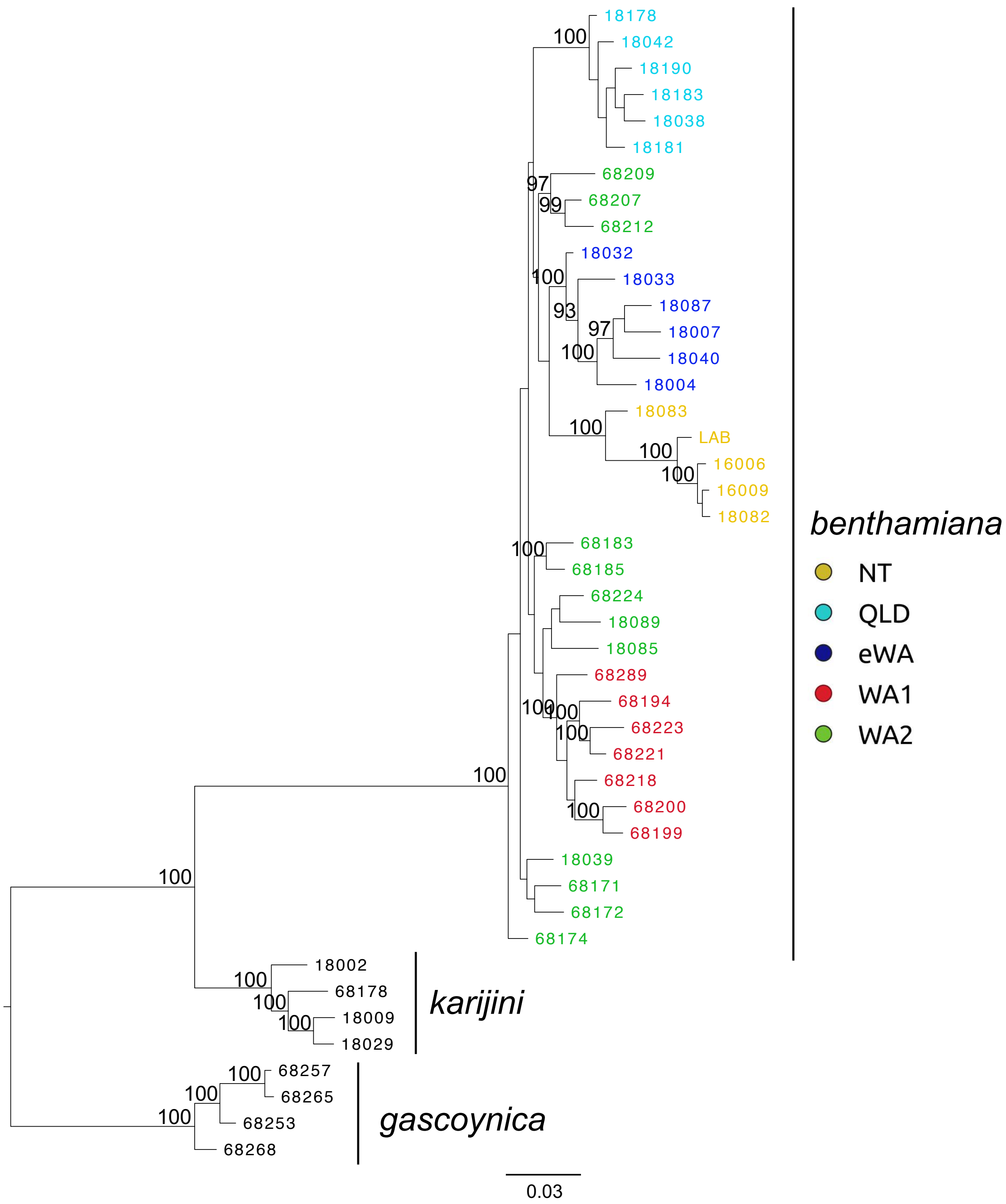


Supplementary Fig. 2. RaxML tree based on 14,285 SNPs. The colour of clusters indicates their geographical provenance: yellow – Northern Territory and northeastern-most Western Australia (NT); dark blue – the deserts of eastern Western Australia (EWA); light blue – Queensland and western-most northern Territory (QLD); red - Pilbara coast of northwestern Western Australia (WA1) and; green - the Pilbara Craton of Western Australia (WA2), the last unresolved but the interrelations of these accessions and their relationships to the other groups are not well supported.

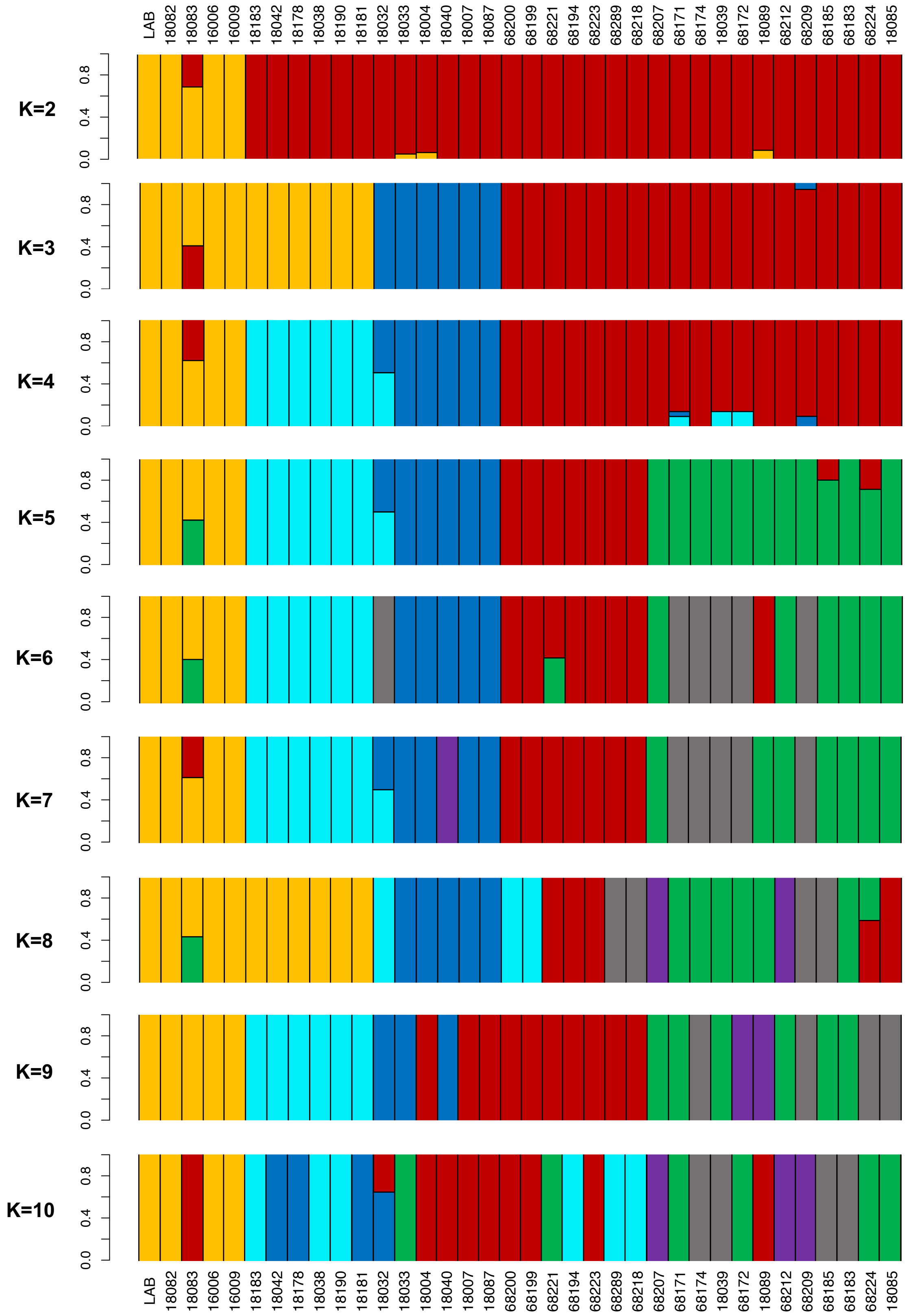


Supplementary Fig. 3. Structure plots of *N. benthamiana* groups obtained in NGSadmix. The values on y-axis represent admixture proportions.

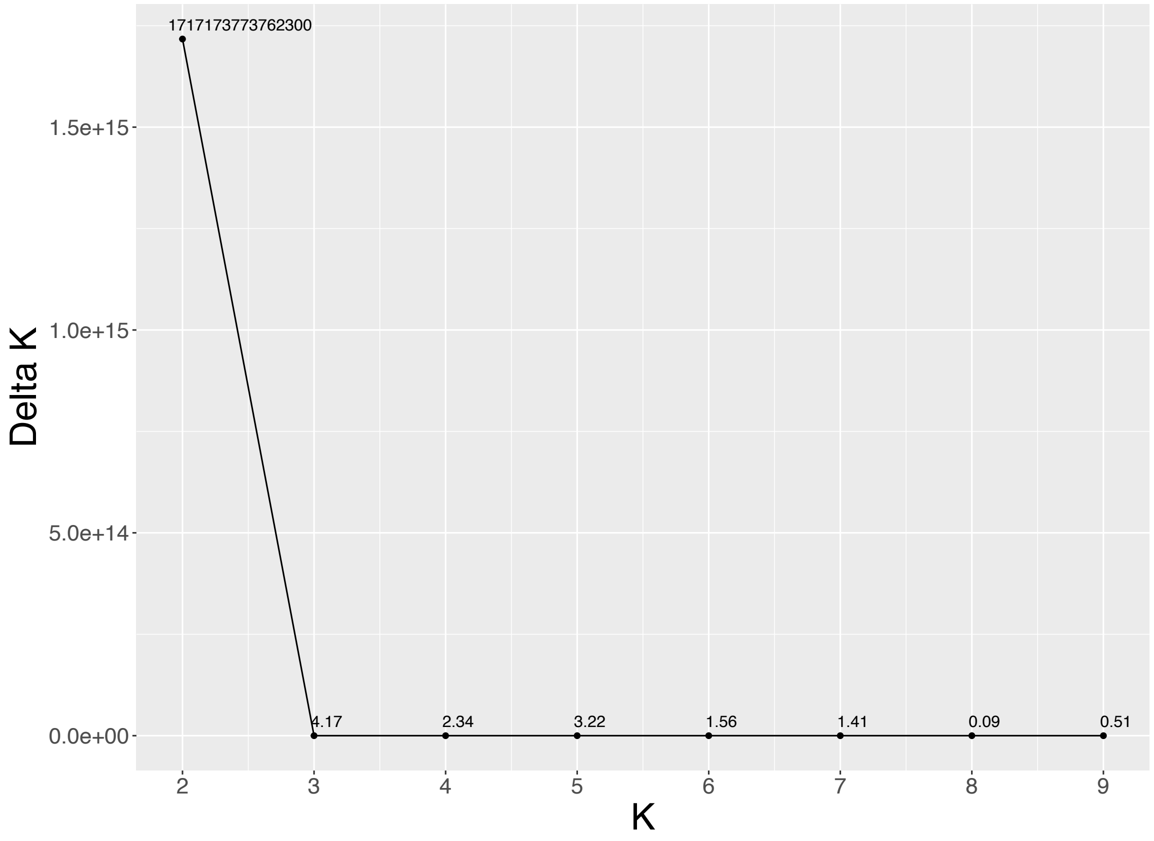


Supplementary Fig. 4. Delta K values for the best K model obtained in NGSadmix. The best K was estimated in accordance with the Evanno method (http://clumpak.tau.ac.il/bestK.html).

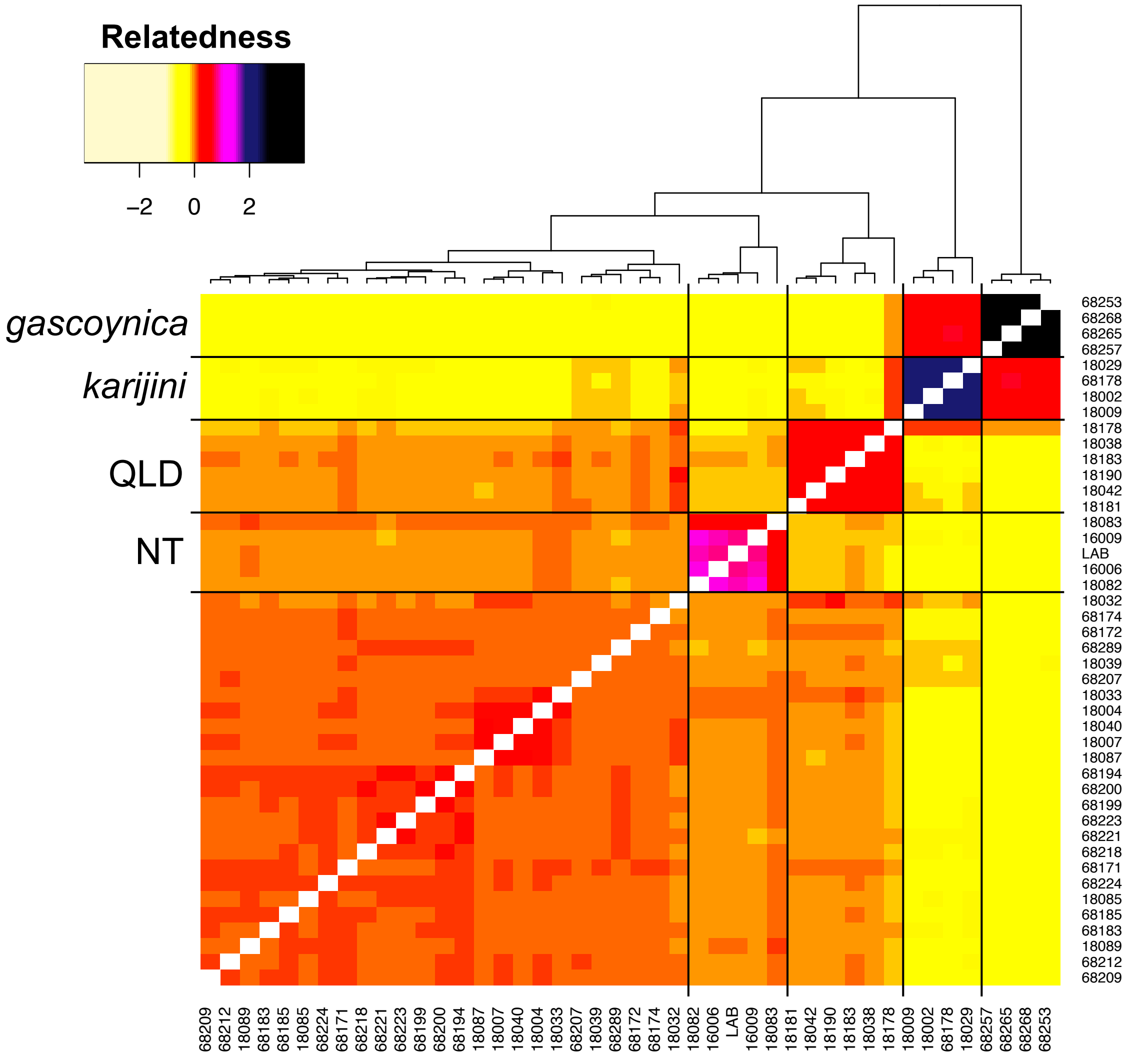


Supplementary Fig. 5. Coancestry heatmap of *N. benthamiana* and the closest related species *N. gascoynica* and *N. karijini*. The heatmap was constructed based on genotype likelihoods obtained in ANGSD. Darker tones represent higher pairwise relatedness according to legend; estimates for the relationship of one individual to itself have been excluded.

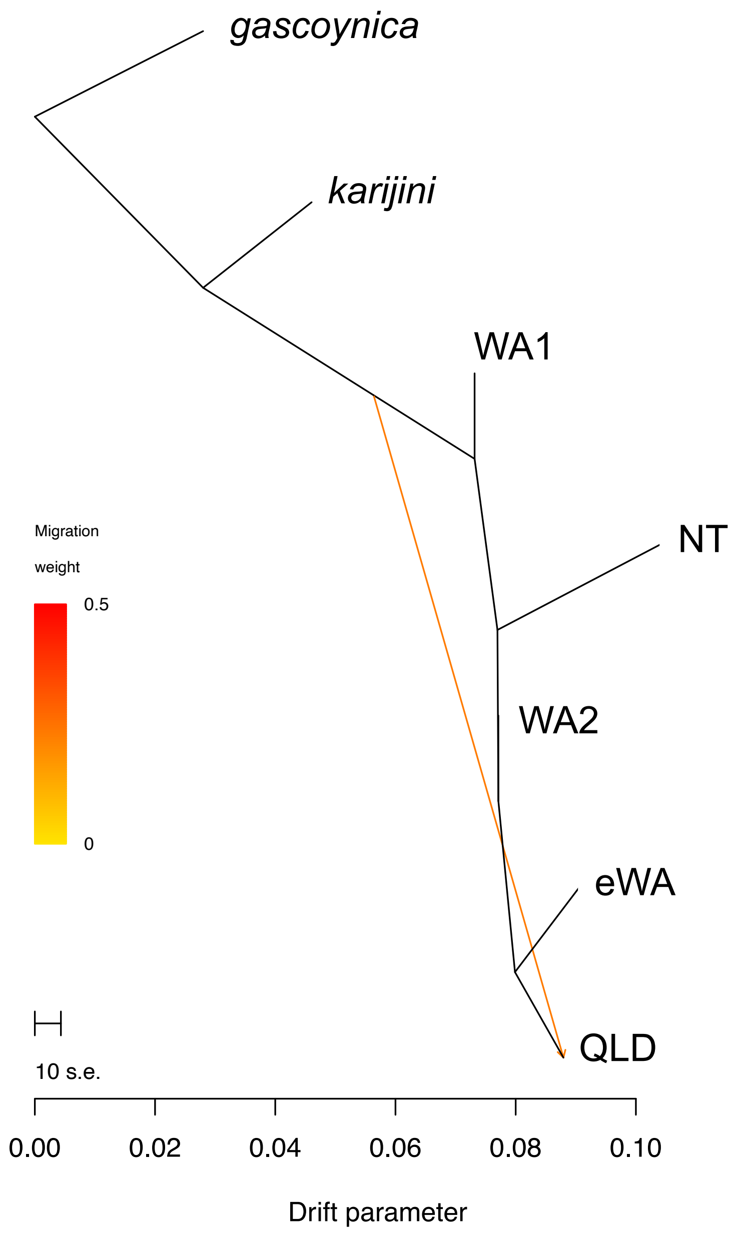


Supplementary Fig. 6. Maximum-likelihood tree produced by TreeMix showing a migration event (arrow) between *N. karijini* and the *QLD* accessions.
